# Efficient cell-free translation from diverse human cell types

**DOI:** 10.1101/2024.12.16.628677

**Authors:** Jana Ziegelmüller, Nikolaos Kouvelas, Priyanka Thambythurai, Alexander M. Hofer, Oliver Mühlemann, Evangelos D. Karousis

## Abstract

The difficulties in producing cell-free translation systems from different cell types limit the ability to study regulatory mechanisms that depend on different biological contexts. Developing systems tailored to diverse cell types would be instrumental in investigating cell-type-specific translational control, co- and post-translational modifications, and viral manipulation strategies. Our method addresses this gap by providing a scalable and adaptable solution for producing high-quality lysates that reflect the specific needs of different cell types.

**Summary:** Cell-free translation systems are indispensable for studying protein synthesis, enabling researchers to explore translational regulation across different cell types. Here, we present a scalable method for preparing translation-competent lysates from a range of frequently used human cell lines using dual centrifugation. We optimized lysis conditions for adherent and suspension cells, producing high-quality lysates from HEK-293 (adherent and in suspension), HeLa, SH-SY5Y, and U2OS cells. Our results demonstrate that cell-specific factors influence translation efficiency, with adherent HeLa cells showing the highest activity. We also observed that sensitivity to lysis conditions varies between cell lines, underscoring the importance of fine-tuning parameters for efficient protein production. Our method provides a robust and adaptable approach for generating cell-type-specific lysates, broadening the application of *in vitro* translation systems in studying translational mechanisms.

## Introduction

Cell-free biology describes the recapitulation, study, and exploitation of complex biological processes without intact cells (1) and has facilitated major advances in diverse research fields by enabling detailed mechanistic explorations and overcoming limitations owing to the complexity of intact cells. In the past, cell-free or *in vitro* translation has contributed to key discoveries, such as the elucidation of the genetic code (2). Recent improvements in cell-free translation systems have expanded their capabilities and potential applications in synthetic biology, biotechnology, and biomedicine. Some recent examples include findings in the features of aminoacyl tRNA synthetases and tRNAs (3), alternative ways to produce proteinosomes, particles that support cell-free transcription and translation (4), and the study of riboswitches (5). Cell-free expression systems have also been more systematically utilized to study the folding of membrane proteins, which are often challenging to express in living cells (6). Furthermore, constructing artificial cells based on cell-free systems (7), integrating cell-free expression reactions in artificial biomolecular condensates (8), and inhibiting translation in synthetic cells using membrane-less organelles (9) fuel the potential for future advances in synthetic biology.

*In vitro* translation systems based on cell lysates allow for the rapid and efficient synthesis of proteins in a cell-free environment, enabling researchers to study protein translation without needing intact cells. The Rabbit Reticulocyte Lysate (RRL) is a cell-free, lysate-based system derived from the immature red blood cells (reticulocytes) of rabbits that is widely used in molecular biology and biochemistry for *in vitro* protein synthesis (10). This system provides a rich source of the components necessary for translation, including ribosomes, tRNAs, amino acids, and various translation factors. However, RRL is of limited use for studying the mechanism of mammalian translation. The ability to translate mRNAs without a 7-methyl-G cap structure and cap-binding proteins and its non-canonical translation initiation activity at internal mRNA sites limit the physiological relevance of this system (11). High levels of globin mRNA and tissue-specific RNases can interfere with translation and degrade mRNA, reducing both efficiency and yield (10,11). Therefore, the specialized nature of reticulocytes may not accurately represent translation processes in other cell types. Additionally, RRL cannot be easily genetically manipulated to deplete or enrich specific protein factors (2). These factors can affect the specificity, control, and broader applicability of the RRL system in certain experimental contexts (2).

Given the limitations of RRL and the ethical issues associated with its production, there has been a growing interest in developing cell-free translation systems based on lysates from immortalized human cells. Such systems offer a more physiologically relevant environment, capturing tissue- and cell-type-specific factors critical for proper translation in mammalian systems. By preserving human regulatory elements, post-translational modifications, and translation control mechanisms, human cell-derived lysates provide a powerful tool for studying translation in a context resembling *in vivo* conditions. This has significant implications for fields like synthetic biology, structural biology, and drug screening, where accurate recapitulation of human cellular processes is essential (2, 14).

Efficient cell-free translation systems have been produced from various mammalian cell lines and used to study the mechanism of translation and translation-relevant processes (15–18). A common challenge of these protocols is the variability in translation efficiency depending on the lysis conditions, leading to significant batch-to-batch variation. To address these problems, we previously developed an *in vitro* translation system based on HeLa S3 suspension cells (19). The key innovation of our approach was to lyse the cells by dual centrifugation, a technique traditionally used in the chemical industry that provides efficient sample homogenization by applying well-controlled shearing forces. We established dual centrifugation as a robust method for routine cell lysis under various buffer conditions, yielding lysates capable of cap- and IRES-dependent translation. The efficacy of this new method is documented in studies exploring the role of Nsp1, a virulent factor from SARS-CoV-2, in human cells (20–22), and translation re-initiation (23).

Here, we report the production of translation-competent lysates from different commonly used human cell lines using our dual centrifugation approach. We used adherent HEK-293 cells to benchmark the production of translation-competent lysates from adherent cells, and we adapted the protocol to produce lysates for cell-free translation from adherent HeLa, U2OS, and SH-SY5Y cells, as well as from HEK FreeStyle 293 suspension cells, from here on referred to as HEK FreeStyle cells.

## Results and discussion

### Optimized conditions to yield translation-competent lysates from adherent HEK-293 cells

Previously, we streamlined the production of translation-competent lysates using HeLa S3 suspension cells by dual centrifugation (DC) and have successfully used it to study translation-related processes (19). Acknowledging the importance of examining molecular mechanisms across different cell types, we set out to test this method’s ability to produce translation-competent lysate from HEK293 Flp-In T-Rex cells, from here on referred to as HEK FITR cells. For this purpose, we grew the cells to 80-90% confluency, harvested them by trypsinization, and resuspended them in a translation buffer supplemented with essential salts and an energy regeneration system consisting of creatine phosphate and creatine kinase, previously optimized for HeLa S3 *in vitro* translation (19). We then subjected the cell suspensions to a range of DC conditions, with speeds varying from 500 to 1’200 rpm for 1 to 4 minutes. The processed lysates, namely the supernatant after spinning down the DC-lysed material, were used to translate a 3xFLAG tagged Renilla luciferase (3xFLAG-RLuc) reporter mRNA for 1 hour at 37°C, followed by luminescence measurements (Fig. 1A). Even though we obtained translation-competent lysates from almost all conditions, translation efficiency depended on both the duration and intensity of DC treatment, with highest RLuc signals observed at 800 and 900 rpm for 1 minute. These conditions differ from the DC conditions for HeLa S3 lysate, which yield best translation activity at 500 rpm for 4 minutes (19), suggesting that different cell types require different lysis conditions to achieve optimal translation efficiency of the derived lysates. To monitor the cell lysis efficiency under the different DC conditions, we performed Trypan Blue staining, where living cells are visible as white spheres, whereas dead cells and nuclei as dark blue foci. As shown in Fig. 1B, the lysis efficiency depends on the applied centrifugal forces. Similar to our experience with HeLa S3 cells, at the optimal condition (800-900 rpm for 1 minute), approximately 50% of the cells were lysed with the cell nuclei remaining intact. DC at higher rpm or for longer periods leads to a higher proportion of lysed cells but concomitantly lower translation efficiency of the resulting lysates. This may result from damaging sensitive components essential for translation or disruption of cellular organelles such as the nucleus, which could release nuclear components that negatively impact translation.

**Figure 1.**
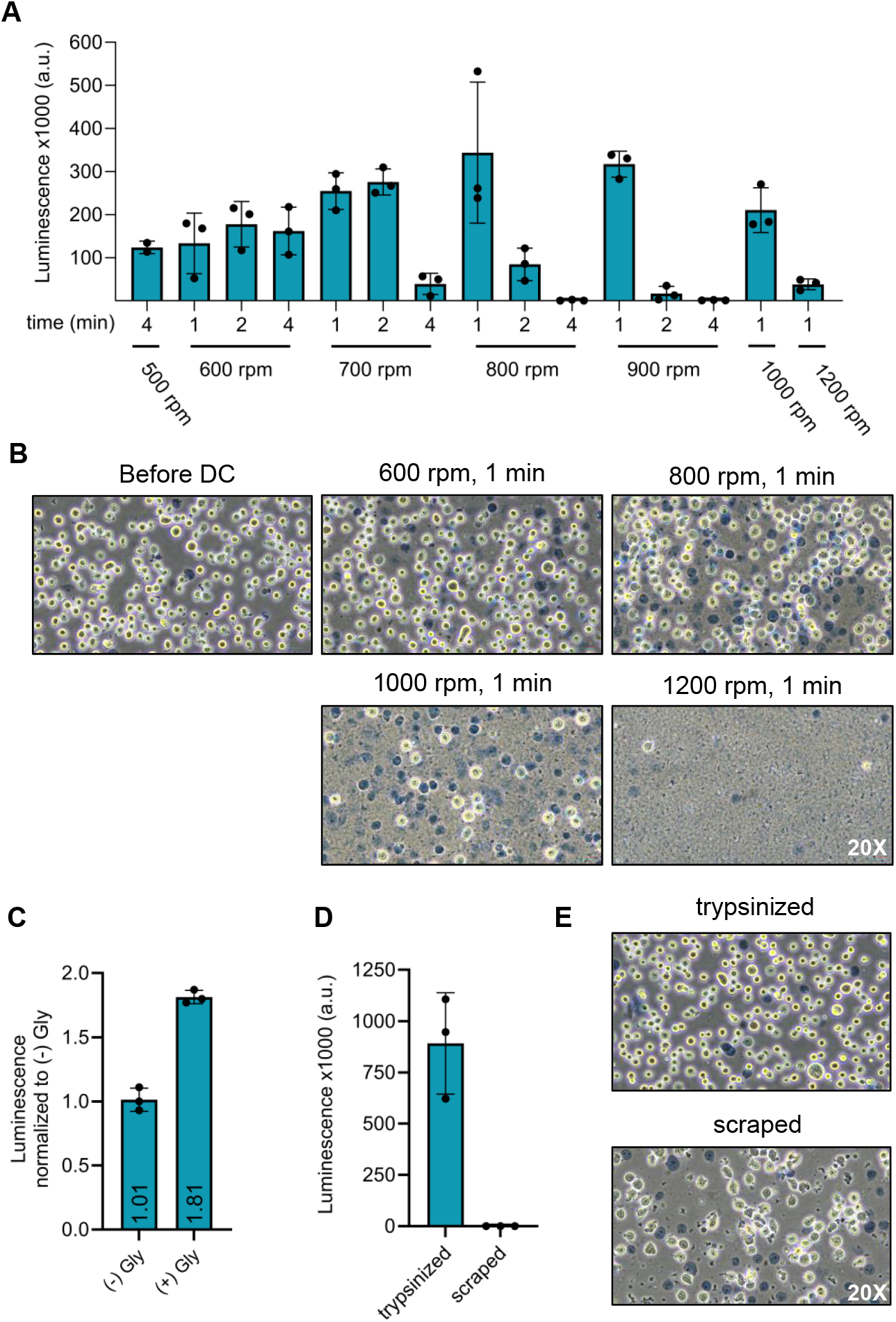
Preparation of translation-competent lysate from HEK FITR cells using DC. **(A)** Comparison of luminescence output HEK FITR lysates produced under different dual centrifugation (DC) conditions by Renilla luciferase assay. **(B)** Examination of HEK FITR cell integrity after DC by Trypan Blue staining. **(C)** Luciferase assay comparing *in vitro* translation efficiency of HEK FITR lysates prepared with or without 15% glycerol in the translation buffer. **(D)** Comparison of translation efficiency between lysates from HEK FITR cells harvested by trypsinization or scraping. The cells were resuspended in a translation buffer containing 15% glycerol and lysed by DC (800 rpm, 1 min, -5°C) after harvesting. **(E)** Trypan Blue staining of HEK FITR cells harvested by trypsinization (top) or scraping (bottom). In (A, C and D) each dot depicts the value of an individual experiment for which the luminescence was measured three times. Mean and SD are shown. All translation reactions were performed using a lysate concentration of 1×10^5^ cell equivalents/ μl containing 5 fmol/ μl 3xFLAG-RLuc mRNA, at 37°C for 1h in a total volume of 25 μl of which everything was used for the Renilla luciferase assay.

Attempting to further improve the translation efficiency of the HEK FITR lysate, we assessed the effect of glycerol on cell-free translation based on its molecular crowding effect (14). We observed an 80% increase in translation efficiency upon adding 15% glycerol to the translation buffer in which the cells are suspended and lysed (Fig. 1C). Notably, addition of glycerol did not increase the translation efficiency of HeLa S3 lysates, suggesting that these lysates may already translate with their maximal translation capacity (Supplementary Fig. 1A). In contrast, HEK FITR lysates might be less concentrated and, therefore, benefit from increasing the local concentration of translation factors by adding a crowding agent.

To speed up the lysate preparation procedure, we further tested whether we could harvest the cells by scraping of the plates instead of the more time-consuming trypsinization procedure. However, we observed that only lysates originating from trypsinized cells were translationally competent (Fig. 1D). Harvesting the cells by scraping is a harsh treatment that damages and lyses a large proportion of the HEK FITR cells, which was clearly visible when the cells were microscopically examined directly after harvesting, before DC (Fig. 1E). Since scraping can influence the integrity of cellular membranes (24), we speculate that the resulting release of inhibitory factors from membrane-bound compartments might be responsible for the reduced translational activity. Alternatively, the mechanical stress of scraping may inactivate critical elements of the translation machinery that are preserved when cells are trypsinized.

In summary, our optimal lysate preparation protocol for HEK FITR cells includes harvesting the cells by trypsinization and resuspension after washing in a translation buffer containing 15% glycerol and DC at 800 rpm for 1 minute.

### Characterization of HEK FITR translation-competent lysate

To characterize the HEK FITR lysate, we first tested the duration of its translation activity using the 3xFLAG-RLuc reporter mRNA. We found that protein production increases linearly for at least one hour of incubation at 37°C (Fig. 2A, B). Furthermore, we titrated the reporter mRNA concentration in the translation reaction and observed an increase in translation at a concentration range of 2.5-20 fmol of reporter mRNA per μl of cell-free translation reaction (Fig. 2C).

**Figure 2.**
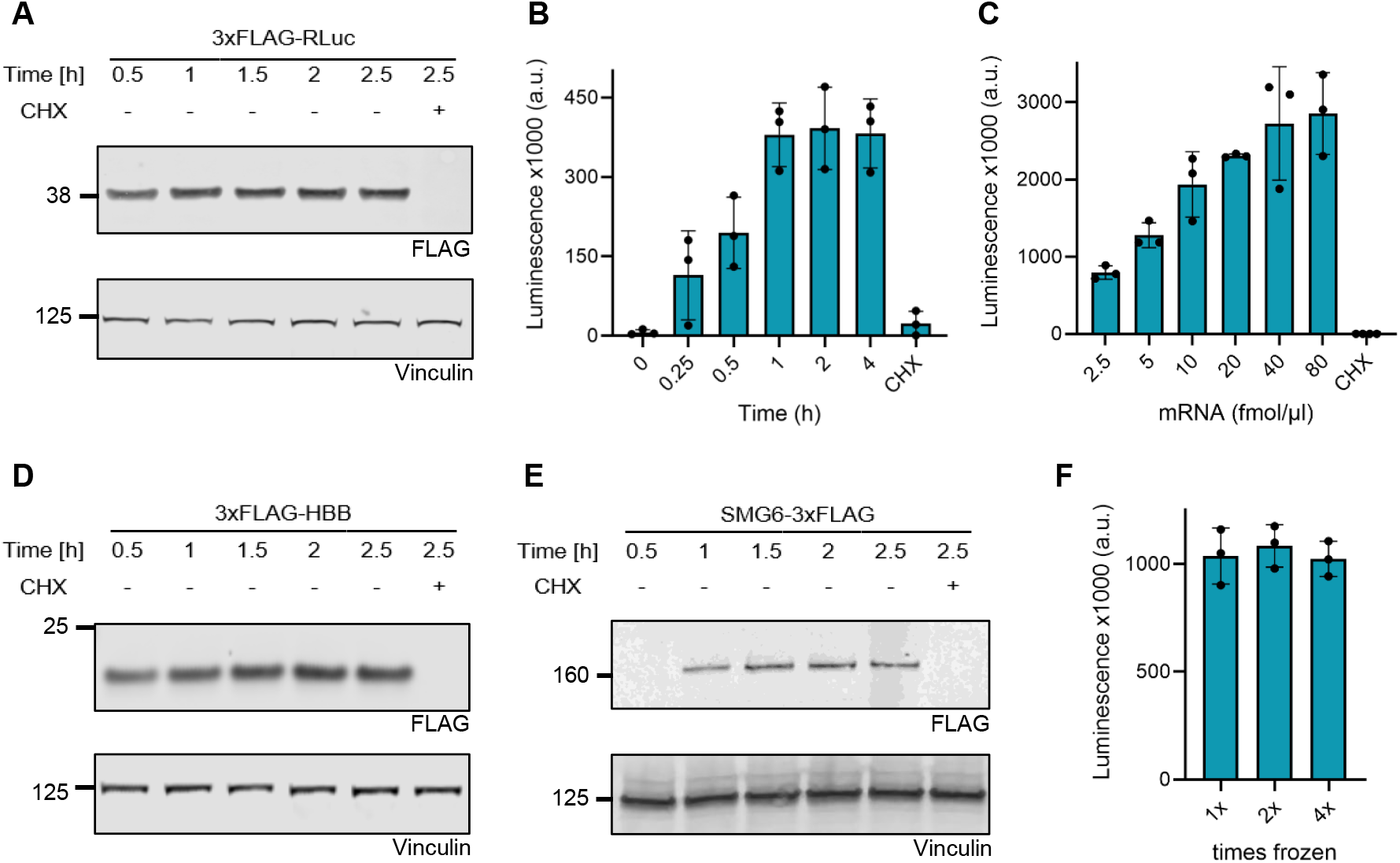
Characterization of HEK FITR lysate. **(A-B)** Western blot analysis and Renilla luciferase assay of time-course *in vitro* translation reactions using the 3xFLAG-RLuc mRNA reporter. **(C)** Renilla luciferase assays of *in vitro* translation reactions with varying 3xFLAG-RLuc reporter mRNA concentrations. **(D-E)** Western blot analysis of time-course i*n vitro* translation reactions using 3xFLAG-HBB and SMG6-3xFLAG mRNAs, respectively. **(F)** Renilla luciferase activity measurements of *in vitro* translation reactions of 3xFLAG-RLuc in HEK FITR lysate subjected to snap-freezing for 1x, 2x or 4x. For (B, C and F), translation reactions with a total volume of 12.5 μl were prepared and were used for the luciferase analysis. Each dot depicts the value of an individual experiment for which the luminescence was measured three times. Mean and SD are shown. For (A, D and E), translation reactions with a volume of 25 μl were performed, of which 2.8 μl were loaded on the gel. Antibodies used are depicted at the bottom of the blots with vinculin serving as a loading control. All translation reactions were performed with a lysate concentration of 1×10^5^ cell equivalents/ μl and 5 fmol RNA/ μl translation reaction at 37°C for 1h. Translation was inhibited in control samples by adding 0.1 mM cycloheximide (CHX).

To assess the versatility of the HEK FITR lysate, we explored the translation of reporter mRNAs encoding proteins of different sizes, including human beta-globin (HBB) and SMG6, an endonuclease involved in nonsense-mediated mRNA decay (NMD) (25, 26). Western blot analysis revealed that consistent with our findings for 3xFLAG-RLuc (37 kDa), we detected distinct single bands for 3xFLAG-HBB (21 kDa) and SMG6-3xFLAG (163 kDa) (Fig. 2A, D, E), demonstrating the HEK FITR lysate’s ability to translate efficiently diverse proteins up to a size of at least 160 kDa and probably beyond, comparable to our previous results with HeLa S3 lysates (19).

To evaluate the stability of the HEK FITR lysate, we tested its resistance to freeze-thaw cycles. As shown in Fig. 2F, the translation output of HEK FITR lysate remains unchanged through at least four freeze-thaw cycles, providing significant storage and handling advantages over other lysates such as RRL, which is recommended to only be used for up to two freeze-thaw cycles (10).

### Production of translation-competent lysates from various cell types

Using translation-competent lysates from different cell types facilitates studying cell-specific differences in translation, revealing tissue-specific regulatory mechanisms and responses. To this end, we addressed to what extent our protocol can be adapted to produce translation-competent lysates from different cell lines. Having succeeded in producing translation-competent lysates from HeLa S3 suspension cells and adherent HEK FITR cells, we next attempted to produce lysates from HEK FreeStyle, a HEK 293-based suspension cell line, and from adherent HeLa cells. To expand the range of cell-type-specific translation-competent lysates, we complemented our panel with two additional commonly used cell lines, SH-SY5Y and U2OS cells, which are of neuroblastoma and osteosarcoma origin, respectively.

Our previous work on HeLa S3 cells (19) and HEK-FITR cells (Fig.1) highlighted the importance of optimizing the cell lysis parameters to yield lysates with high translational activity.

Therefore, we tested different DC parameters for each cell line, beginning with the optimal conditions for HeLa S3 and HEK-FITR cells (500 rpm for 4 min and 800 rpm for 1 min). We then increased or decreased the DC rpms and duration, based on the translation efficiency of the initial lysates, until we identified conditions that maximized translation of the 3xFLAG-RLuc reporter protein (Fig. 3). Furthermore, we monitored cell lysis efficiency by Trypan Blue staining of the cell suspension samples after DC treatment (Supplementary Fig. 2).

**Figure 3.**
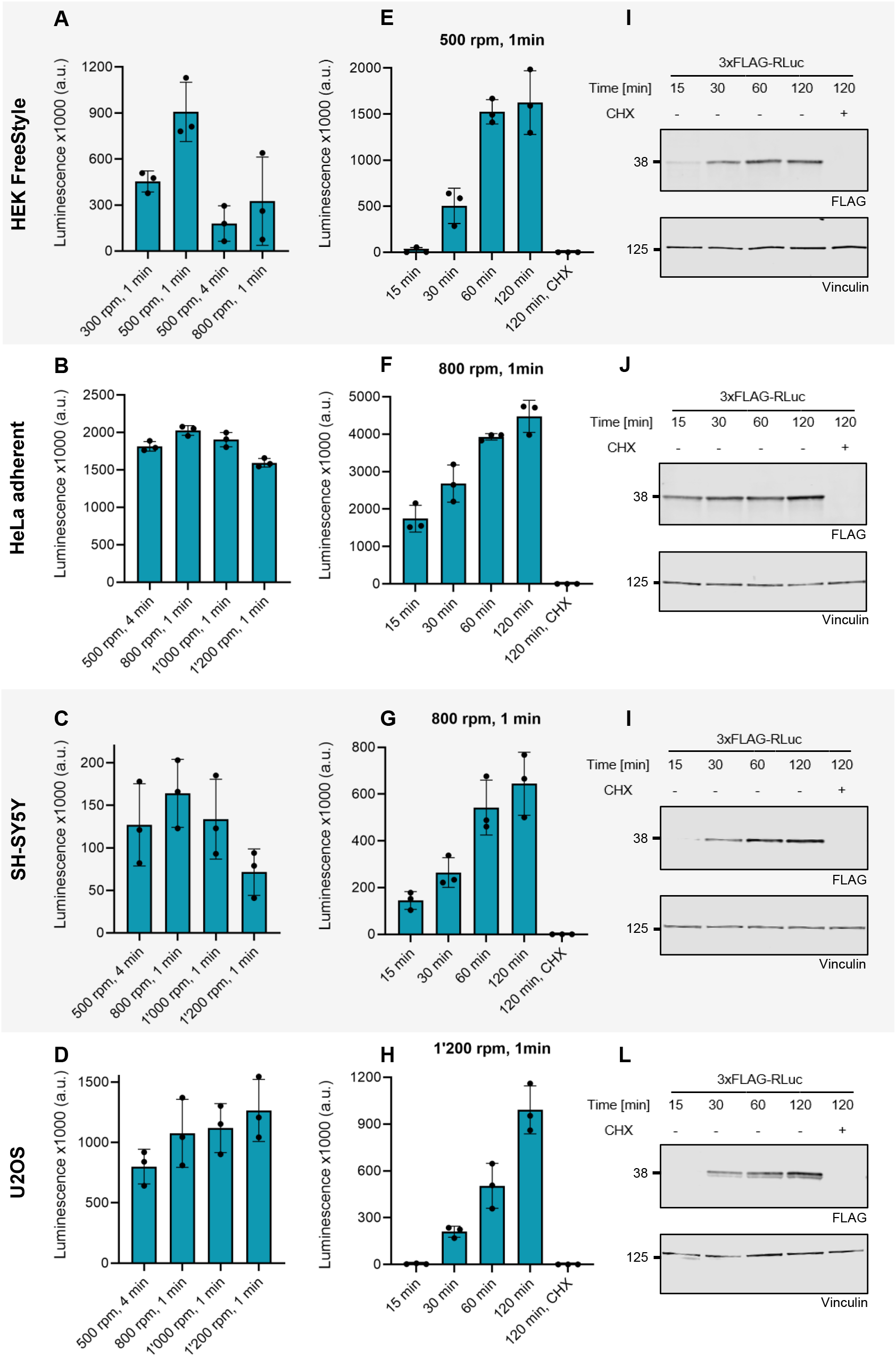
Production of translation-competent lysate from various cell lines. **(A-D)** Optimization of dual centrifugation (DC) conditions for lysate preparation from HEK FreeStyle, HeLa adherent, SH-SY5Y and U2OS cells, respectively. Translation efficiency was assessed by luciferase assay performed following *in vitro* translation of the 3xFLAG-RLuc reporter mRNA for 1h at 37°C. The DC conditions tested are indicated on the y-axis. **(E-H)** Renilla luciferase assay of time-course *in vitro* translation of the 3xFLAG mRNA in lysates of the different cell lines. The lysates employed were prepared under the conditions that yielded the highest translation efficiency (see A-D), with these optimal conditions noted at the top of each graph. **(I-L)** Western blot analysis of translation reactions from (E-H). The antibodies used are depicted below each blot, with vinculin serving as a loading control. In (A-H), each dot depicts the value of an individual experiment for which the luminescence was measured three times. Mean and SD are shown. For (A-D) and (E-L), 25 and 40 μl translation reactions were performed, respectively. For (A-H) 25 μl of the translation reactions were used for luciferase assay. For (I-L) 2.8 μl of the translation reactions were loaded on the gel. All translation reactions were performed with a lysate concentration of 1 x10^5^ cell equivalent / μl and 5 fmol 3xFLAG-RLuc mRNA/ μl translation reaction. 0.1 mM cycloheximide (CHX) was used to inhibit translation in control samples.

Both, the translation efficiency readout and the Trypan Blue staining (Fig. 3A-D and Supplementary Fig. 2) showed that certain cell lines, such as HEK FreeStyle, are more sensitive to DC and minor alterations of the lysis parameters, while others, like adherent HeLa cells, resulted in similar degrees of lysis and translation efficiency over a wide range of DC conditions. Interestingly, U2OS cells displayed a notably high lysis efficiency at the optimal condition. The observed differences in cell sensitivity to lysis parameters may be attributed to the distinct structural characteristics of these cell types.

Next, we assessed how the duration of *in vitro* translation is associated with producing the 3xFLAG-RLuc reporter protein. For this purpose, we conducted time-course translation experiments in the lysates produced from the different cell lines under their above-determined optimal DC conditions. Translation efficiency was assessed based on 3xFLAG-RLuc using luminescence and Western Blot measurements. As shown in Fig. 3E-L, we observed an increase in Renilla luciferase activity during the first hour of translation, which reached a plateau at two hours in all cases, apart from the U2OS-derived lysate, which translated for a longer time. Thus, the reporter protein output increased over a shorter time window for all the newly tested cell lines, except for U2OS, compared to HeLa S3 lysates. As reported previously (19), HeLa S3 lysates increased the reporter protein output for at least 2 hours of translation reaction (Sup. Fig. 1B).

Successful production of translation-competent lysates from diverse cell lines demonstrates the adaptability of the protocol, broadening its application for studying cell-type-specific translation mechanisms. The different sensitivity of different cell types to lysis conditions underscores the need to optimize cell lysis protocols based on the unique structural characteristics of each cell type. This variation may impact lysate quality and downstream applications, highlighting the importance of fine-tuning dual centrifugation parameters. The plateau in translation efficiency over time indicates possible limitations in translation machinery or substrate availability over time, which may vary across cell types, necessitating further investigation into rate-limiting factors.

Having optimized the lysate preparation protocol for each cell line, we next compared the translation efficiency of the different cell type-specific lysates after *in vitro* translation for 1 hour. The translation efficiency, measured through 3xFLAG-RLuc reporter activity and Western Blot analysis, shows a marked difference between cell lines (Fig. 4A, B). Notably, the adherent HeLa cell lysate exhibited the highest translation efficiency, while SH-SY5Y and U2OS lysates demonstrated considerably lower levels of translation. Both HEK 293 lysates, from adherent FITR and suspension FreeStyle cells, showed similar translation efficiencies. It is important to note that these results reflect only a snapshot of translation efficiency for the different lysates under this specific translation condition. Adjusting the translation parameters could yield different outcomes. For example, extending the translation duration from 1 hour to 2 hours would result in considerably higher translation efficiencies in both U2OS and HeLa S3 lysates (Fig. 3H and L, Supplementary Fig. 1B).

**Figure 4.**
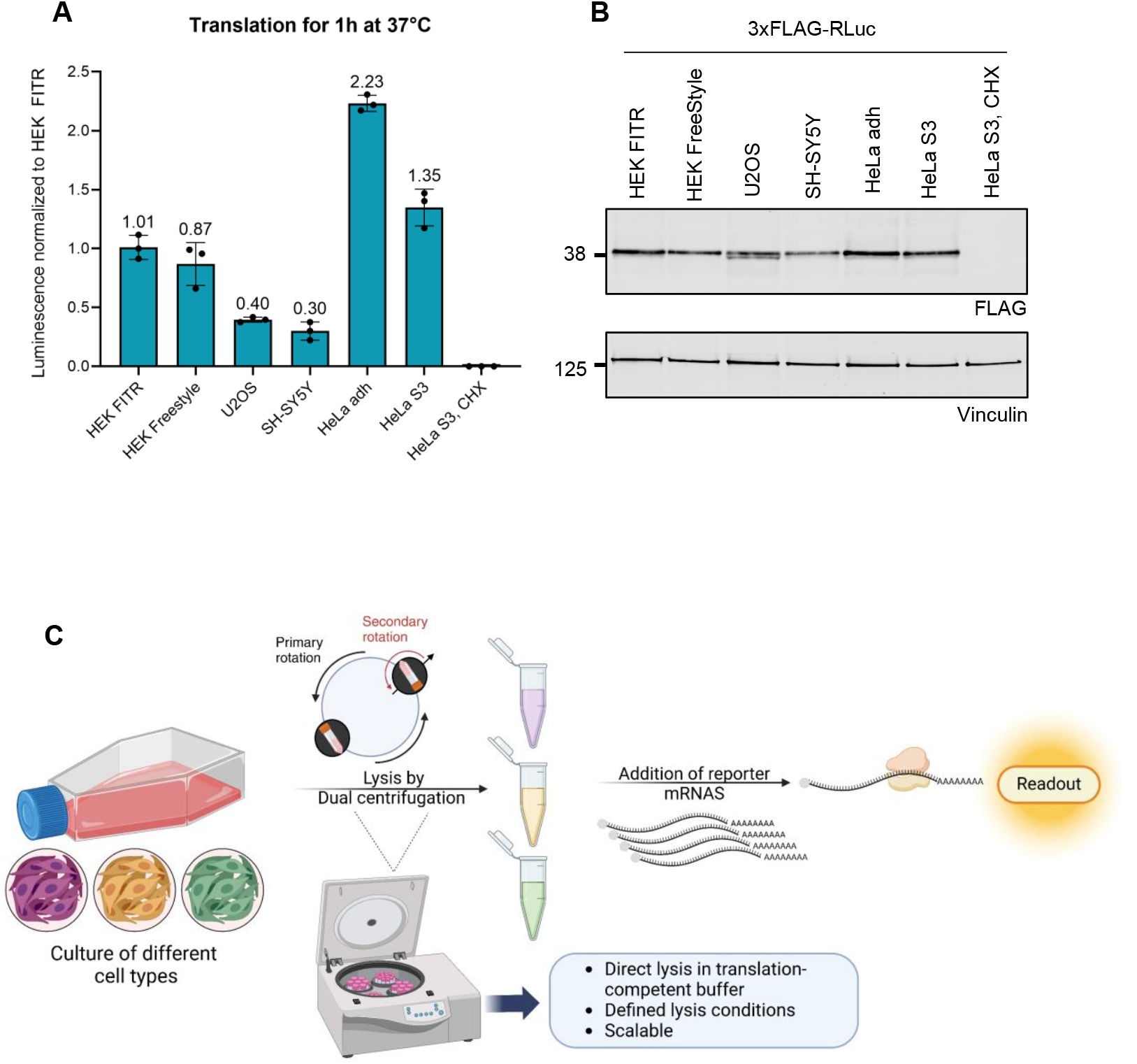
Comparison of translation efficiency of lysates from different cell types. **(A)** Renilla luciferase assay comparing the translation efficiency of the different lysates presented in this study. Each dot depicts the value of an individual experiment for which the luminescence was measured three times. Mean and SD are shown. **(B)** Western blot analysis of the corresponding translation reactions from (A). The antibodies used are depicted below the blots, with vinculin serving as a loading control. For (A-B) *in vitro* translation reactions contained 5 fmol 3xFLAG-RLuc mRNA/ μl and were performed at 37°C for 1h with a lysate concentration of 1×10^5^ cell equivalent/ μl in a total volume of 40 μl. 0.1 mM cycloheximide (CHX) was used to inhibit translation in control samples. 15 μl of the reactions were used for the Renilla luciferase assay, and 2.3 μl were loaded on the gel for Western blot analysis. **(C)** Schematic depiction of the workflow for preparing translation-competent lysate from various cell types created with Biorender.com.

The variation in translation efficiency across cell lines could reflect the importance of cell-type-specific factors in regulating translation but might also indicate different degrees of cell lysis or reduced activity of translation factors in these cell types.

Overall, we adapted DC parameters, as illustrated in Figure 4C, to achieve efficient lysis and produce translation-competent lysates. This method demonstrates versatility across various cell types, allowing us to adapt the conditions for both adherent and suspension cells. Notably, the scalability of this approach makes it suitable for diverse cell-type-specific translation studies ranging from small-scale functional studies to large-scale biochemical purification experiments.

The possibility of producing translation-competent lysates from specific cell lines is instrumental for several reasons. Many translation-relevant processes require tissue-specific factors, and some proteins only acquire post-translational modifications in specific cell-type derived lysates and not in others. For instance, extracts from HeLa cells proved inefficient in producing an N-glycosylated form of human immunodeficiency virus type-1 envelope protein 120 (gp120), while the hybridoma extract was able to fully N-glycosylate gp120 (27).

The ability to generate lysates from various cell cultures creates new opportunities to produce lysates from genetically modified cells, allowing for controlled variations in factors of interest through knockdown, knockout, and overexpression experiments. The protein production output can be further increased by the addition of auxiliary proteins and other modifications of parameters that can increase the translation rate, as previously reported (19, 28).

The success of the DC-based cell lysis protocol across different cell lines highlights the versatility and adaptability of the method. We anticipate our lysate preparation can be applied to virtually any cell line, facilitating studies of cell-type-specific translation mechanisms across a wide range of biological and disease contexts. This opens exciting possibilities for both basic research and industrial applications, enabling the production of cell-free translation systems tailored to specific cellular environments. The ability to scale this process and produce ample amounts of translation-competent lysates renders it a valuable tool for high-throughput applications and facilitates cross-cell-type comparisons of translational activities. Furthermore, it is well suited for isolating cell-type-specific translation complexes in sufficient amounts and purity for cryogenic electron microscopy studies.

Cell-free translation systems are instrumental in studying translational mechanisms and producing proteins of interest in various biological contexts. Our method expands the possibility of producing translation-competent lysates from specific cell types, providing systems that are promising for the investigation of co- and post-translational phenomena, such as cell-type specific translation phenotypes (29), post-translational modifications (27), and viral manipulation (30, 31), which can differ between cell types.

### Limitations of the study

While the DC method for preparing translation-competent lysates has proven versatile across various cell lines, each new cell type-requires careful optimization of the lysis conditions, which is determined mainly by the DC speed and time. Achieving the right balance between disruption of the cell membrane and preserving translation activity, particularly for more delicate or hard-to-lyse cells, is crucial for maximizing the yield of recombinant protein. Additionally, while the method works well for cultured cell lines, its scalability remains untested for more complex tissues, like primary cells or multi-cellular tissues. These tissues present additional structural complexity, which may impede lysate quality and translation efficiency. Another consideration is the preservation of post-translational modifications, which may not be fully maintained during lysate preparation, potentially affecting studies of cellular regulation. Furthermore, the method may have limitations when applied to studying organelle-specific translation processes, such as those occurring in mitochondria or at the endoplasmic reticulum, where localized translation events play key roles.

## Material and Methods

### General cell culture growth conditions

HEK293 Flip-In T-Rex (HEK FITR), SH-SY5Y and U2OS cells were cultured in DMEM/F-12 (Gibco, Cat No. 32500-043) medium supplemented with 100 U/ml penicillin, 100 μg/ml streptomycin and 10% FCS (BioConcept, Cat No. 2-01F30-I) (termed DMEM+/+), at 37°C and 5% CO_2_ in a humid atmosphere on 150 cm^2^ dishes to 80-90% confluency. HeLa S3 cells were grown in DMEM+/+ medium and were grown for two passages in 150 cm^2^ dishes after thawing before transferring to suspension culture flasks in a volume between 20 ml - 600 ml growing in a range of 0.1×10^6^-1.2×10^6^ cells/ml. HEK FreeStyle 293 cells were cultured in HyClone SFM4HEK293 medium (Cytiva, Cat No. SH30521.02) in volumes between 20 ml – 600 ml in a range of 0.3 × 10^6^ – 2 x10^6^ cells /ml at 37°C and 8% CO_2_ in a humid atmosphere. Suspension culture flasks were used for growing HeLa S3 and HEK FreeStyle 293 cells (Corning flasks Cat No. 431405, 431147, and 431255).

### Translation-competent lysate preparation

To prepare HEK FITR, HeLa adherent, SH-SY5Y, and U2OS lysates, the cells were grown on 8 – 12x 150 cm^2^ cell culture dishes to an 80-90% confluency. To harvest the cells, they were washed with 10 ml PBS, 2 ml trypsin/EDTA was added per plate, and the cells were incubated at 37°C until they were fully detached. When using cell scraping for harvesting HEK FITR cells, the medium was removed, and the cells were scraped in 500 μl PBS. The cells of 4 - 6 dishes were collected in a total volume of 20 ml of DMEM+/+. This step was repeated to collect the cells remaining on the plates. The cells were counted with Trypan Blue staining to calculate the required volume of translation buffer, transferred to 50 ml tubes, and centrifuged at 500 *g* for 2 min at 4°C. The medium was discarded, and all cells were transferred in one 50 ml falcon tube in a total of 30 ml ice-cold 1x PBS pH 7.4. The resuspension was performed gently to prevent premature lysis of the cells. The cells were centrifuged at 500 *g* for 2 min at 4°C, and the supernatant was discarded. The washing step was repeated in 10 ml 1x PBS pH 7.4. Next, the cells were resuspended in ice-cold translation buffer (33.8 mM HEPES pH 7.3, 63 mM K-acetate pH 5.5, 0.68 mM KCl, 13.5 mM creatine phosphate, 230 ng/ml creatine kinase, 1x protease inhibitor cocktail (Bimake, Cat No. B14002), 15% glycerol (unless otherwise stated)) to reach a concentration of 2×10^8^ cells/ml and the suspension was transferred to 2 ml screw cap microtubes (Sarstedt, Cat No. 72693) in 400 μl aliquots per tube. Cell lysis was performed by dual centrifugation using the ZentriMix 280 R system (Hettich AG) at a temperature of -5°C. Cell lysis efficiency was monitored by staining the samples in a 1:6 dilution before and after dual centrifugation with 0.2% Trypan Blue (Thermo Fisher Scientific, Cat No.EVS-050) and observed under a light microscope using EVE Cell counting slides (NanoEnTek, Cat No. EVS-050). The samples were centrifuged at 13’000 *g* for 10 min at 4°C, and the supernatant (lysate) was collected, snap-frozen, and stored at -80°C.

For the production of translation-competent lysate from HEK FreeStyle 293 cells, the cells were grown to a density of 1.8 - 2 × 10^6^ cells /ml in a volume of 200 ml. The cells were counted, and the lysate was prepared as described above. During the washing steps of the cells in PBS, all cells were combined in one 50 ml falcon tube.

HeLa S3 lysate was prepared as described (19) with the modification that the translation buffer (33.8 mM HEPES pH 7.3, 63 mM K-acetate pH 5.5, 0.68 mM KCl, 13.5 mM creatine phosphate, 230ng/ml creatine kinase, 1x protease inhibitor cocktail (Bimake, Cat No. B14002)) utilized in this work was not supplemented with amino acids.

### *In vitro* transcription and capping

Reporter mRNAs were encoded from plasmids with a pCRII backbone. The 3xFLAG-RLuc (humanized) and the 3xFLAG-HBB reporter both contain an AG initiator sequence for capping with the CleanCap reagent AG (TriLink Biotechnologies, Cat No. N-7113-5), the HBB 5’UTR and a short 3’UTR with a length of 20 bp and a BsmFI binding site downstream of the poly(A) sequence for linearization. The SMG6-3xFLAG reporter plasmid is described in (19) and contains the SARS-CoV-2 Leader sequence as the 5’UTR, a 221 bp long 3’UTR, and a *Cla*I cleavage site downstream of the poly(A) sequence. In all reporters, the template-encoded poly(A) sequence is 33 nts long.

Before *in vitro* transcription, the plasmids were linearized using *BsmF*I (Cat No. R0572L) for 3xFLAG-RLuc and 3xFLAG-HBB and *Hind*III-HF (Cat No. R3104L) for SMG6-3xFLAG in reactions containing 40 ng/ μl DNA (4 μg in total) in 1x CutSmart buffer (NEB, Cat No. B7204S) for 2h or overnight at 37°C, respectively. The linearization was monitored by loading 500 ng DNA on a 1% agarose gel. The linearized DNA was purified using the ChIP DNA Clean & Concentrator kit (ZYMO research, Cat No. D5205), and eluted in 10 μl elution buffer. *In vitro* transcription, using the linearized plasmid as a template (10 μl of elution), was performed in a reaction containing 1x OPTIZYME transcription buffer (Thermo Fisher Scientific, Cat No. BP81161), 1 mM of each NTP (Thermo Fisher Scientific, Cat No. R0481), 1 U/μl RNase inhibitor (Vazyme, Cat No. R301-03), 0.001 U/μl pyrophosphatase (Thermo Fisher Scientific, Cat No. EF0221) and 1.5 U/μl T7 polymerase (Thermo Fisher Scientific, Cat No. EP0111). For the 3xFLAG-RLuc and 3xFLAG-HBB reporter 0.8 mM CleanCap reagent AG (TriLink Biotechnologies, Cat No. N-7113-5) was included in the reaction. The transcription reaction was incubated at 37°C 1h. An additional 1.5 U/μl T7 polymerase was added, and the reaction was further incubated for 1h at 37°C. Subsequently, 0.15 U/ μl Turbo DNase (Invitrogen, Cat No. AM2238) was added, and the reaction was incubated at 37°C for 30 min to digest the plasmid DNA. The transcribed mRNA was isolated using the Monarch RNA Cleanup Kit (New England Biolabs, Cat No. T2040L), eluted in 1 mM sodium citrate pH 6.4 (Gene Link, Cat No. 40–5014-05) and quantified by A_260_ measurement. The 3xFLAG-SMG6 reporter was capped using the Vaccinia Capping System (NEB, Cat No. M2080S) according to the manufacturer’s instructions with the modification that 1 U/ μl RNase inhibitor (Vazyme, Cat. No. R301-03) was added to the reaction. The capped mRNA was isolated and quantified in the same way as after transcription. All *in vitro* transcribed mRNAs were aliquoted, snap-frozen, and stored at -80°C.

### *In vitro* translation

For *in vitro* translation, the lysates were used at a concentration of 1×10^8^ cell equivalents/ ml (stock = 2×10^8^ cell equivalents/ ml) and supplemented with 1 U/ μl RNase inhibitor (Vazyme, Cat. No. R301-03). The *in vitro* transcribed and capped mRNAs were incubated at 65°C for 5 min and cooled on ice before addition, *in vitro* translation was performed at 37°C, and samples were afterwards placed on ice. To inhibit translation as a negative control, 0.1 mM cycloheximide (CHX) was added to selected samples.

### Luciferase assay

Luciferase assays were performed using the Renilla-Glo system (Promega, Cat No. E2720). *In vitro* translation reactions were combined with 1x Renilla-Glo substrate in Renilla-Glo buffer in a 1:4 ratio, transferred to a white-bottom 96-well plate (Greiner, Cat. No. 655073) and incubated at room temperature for 10 min. The luminescence was measured three times using the TECAN infinite M1000 Pro plate reader according to the manufacturer’s guidelines.

### Immunoblot analysis

All samples for western blot analysis were diluted in 1.5x LDS loading buffer (Invitrogen, Cat No. NP0008) containing 50 mM DTT. The samples of 3xFLAG-RLuc and 3xFLAG-HBB were run on mPAGE 4-20% Bis-Tris, 15-well, mini gels (Millipore, mPAGE, 15-well, Cat No. MP42G15) (Figure 2A and D) or Criterion XT 4-12% Bis-Tris, 12+2, midi gels (Bio-Rad, Cat No. 3450123) (Figure 3 and 4). The SMG6-3xFLAG samples were run on NuPAGE 3-8% Tris-Acetate, 10-well, mini gels (Invitrogen, Cat No. EA0378BOX) (Figure 2E). MOPS, MES, and Tris-Acetate buffer were used for 3xFLAG-RLuc, 3xFLAG-HBB, and SMG6-3xFLAG translation reactions, respectively. Proteins were transferred on nitrocellulose membranes (Bio-Rad Cat No. 1704158 (mini) and Cat No. 1704159 (midi)) using the Trans-Blot Turbo Transfer system (Bio-Rad). Subsequently, the membranes were blocked with 5% milk in TBS containing 0.1% tween (TBS-t) and incubated overnight at 4°C with the primary antibodies: Vinculin (Santa Cruz Biotechnology, Cat No. sc-73614, 1:500) and FLAG M2 (Sigma-Aldrich, Cat No. F3165, 1:2’000). The membranes were washed twice with TBS-t for 10 min and incubated for 1h at room temperature with the IR-Dye-conjugated secondary antibodies: IRDye 680LT Donkey anti-Mouse IgG (1:10’000) and IRDye 800CW Goat anti-Mouse IgG2a-specific (1:10’000). Subsequently, the membranes were scanned using the Odyssey Infrared Imaging System 9120 (LiCor).

## Acknowledgements

This work was supported by the National Center of Competence in Research (NCCR) on RNA & Disease funded by the Swiss National Science Foundation (SNSF; grant 51NF40-141735), by SNSF grant 310030-204161 to O.M., by the canton of Bern (University intramural funding to O.M.). Grants awarded to EDK from the Swiss National Science Foundation (SNSF CRSK-3_220624), the Multidisciplinary Center of Infectious Diseases from the University of Bern (MCID), the Fondation Claude et Giuliana, the Forschungsstiftung of the University of Bern, and the Holcim Stiftung. Figure 4C was made using Biorender.

## Author contributions and Declaration of Interests

J.Z., N.K., P.T., and A.H. conducted the experiments; E.K. and J.Z. designed the experiments and E.K. wrote the initial manuscript. All authors read and edited the manuscript. The authors declare no competing interests.

## Supplementary data

**Supplementary Figure 1.**
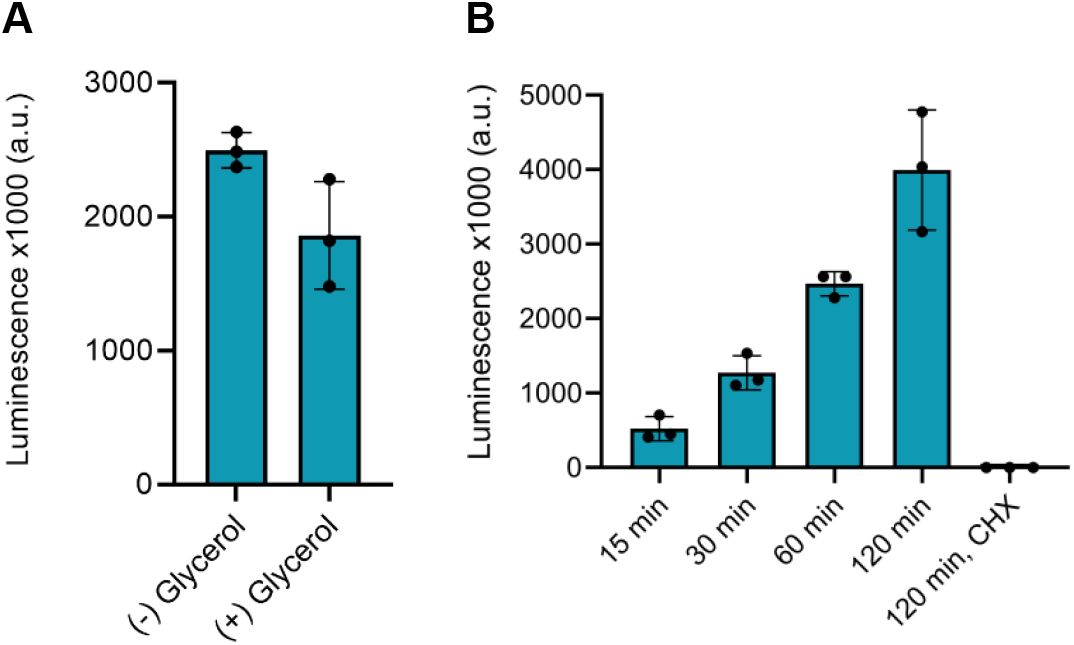
Characterization of HeLa S3 lysate. **(A)** Renilla luciferase assay comparing the translation efficiency of HeLa S3 lysate prepared with or without 15% glycerol in the translation buffer. **(B)** Renilla luciferase assay of time-course *in vitro* translation reactions of HeLa S3 lysate. For (A) and (B) *in vitro* translation reactions contained 5 fmol 3xFLAG-RLuc mRNA/ μl and were performed at 37°C for 1h with a lysate concentration of 1×10^5^ cell equivalent /ml in a total volume of 25 μl of which everything was used for the Renilla Luciferase Assay. Each dot depicts the value of an individual experiment for which the luminescence was measured three times. Mean and SD are shown.

**Supplementary Figure 2.**
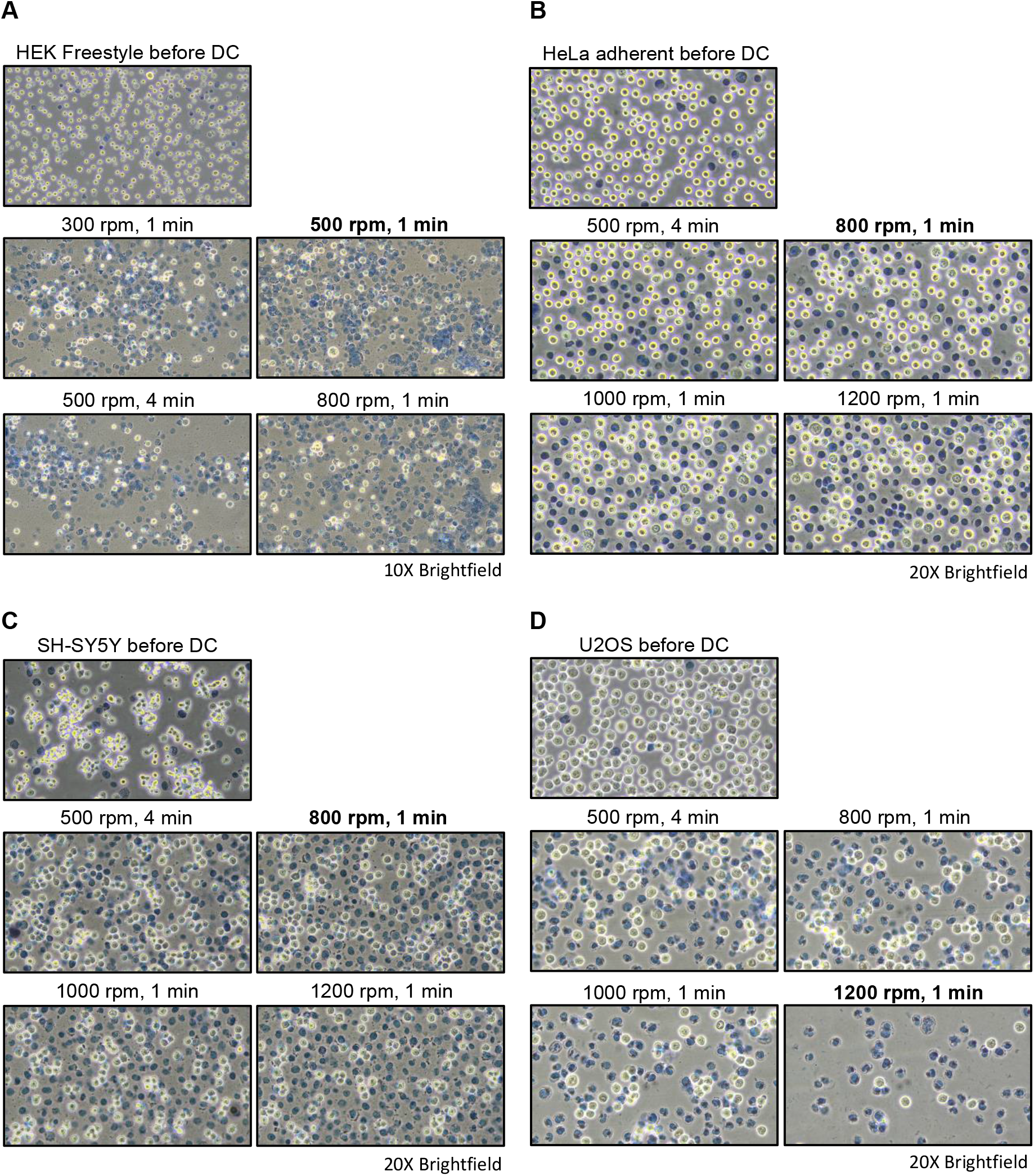
Cell lysis efficiency of different cell lines by dual centrifugation. **(A-D)** Examination of cell integrity of HEK FreeStyle, HeLa adherent, SH-SY5Y and U2OS cells after dual centrifugation assessed by Trypan Blue staining. The corresponding Renilla luciferase assays of the resulting lysates are shown in Figure 3A-D. The DC conditions used are indicated above each image and the condition with the best translation efficiency according to Figure A-D is indicated in bolt letters.

